# Reduced motivation for social contact in Disrupted-in-schizophrenia transgenic rats

**DOI:** 10.1101/2021.09.29.462172

**Authors:** Mohammad Seidisarouei, Marijn van Wingerden, Sandra Schäble, Svenja V. Trossbach, Carsten Korth, Tobias Kalenscher

## Abstract

The Disrupted-in-schizophrenia 1 (DISC1) signaling pathway is considered to play a key role in schizophrenia, depression, autism and other psychiatric disorders. *DISC1* is involved in regulating the dopaminergic neurotransmission in, among others, the mesolimbic reward system. A transgenic rat line tgDISC1 has been introduced as a model system to study behavioral phenotypes associated with abnormal DISC1 pathways. Here, we evaluated the impact of impaired *DISC1* signaling on social (social interaction) and non-social (sucrose) reward preferences in the tgDISC1 animal model. In a plus-maze setting, rats chose between the opportunity for social interaction with an unfamiliar juvenile conspecific (social reward) or drinking sweet solutions with variable sucrose concentrations (non-social reward). tgDISC1 rats differed from wild-type rats in their social, but not in their non-social reward preferences. Specifically, *DISC1* rats showed a lower interest in interaction with the juvenile conspecific, but did not differ from wild-type rats in their preference for higher sucrose concentrations. These results suggest that disruptions of the DISC1 pathway that is associated with altered dopamine transmission in the brain result in selective deficits in social motivation seen in neuropsychiatric illness.

## Introduction

The disrupted-in-schizophrenia 1 (DISC1) protein and its signaling pathway play an important role in mental diseases. DISC1 was originally identified in a Scottish family where chromosomal translocation directly disrupted brain-expressed genes associated with several psychological disorders, such as schizophrenia, bipolar disorder, and recurrent major depression (Millar et al., 2000; Blackwood et al., 2001). Alterations in the DISC1 gene are associated with impairments in brain development in humans, as well as primates and rodents, explicating a possible mechanism for their role in several psychiatric disorders (Austin et al., 2003; Hashimoto et al., 2006; Clapcote et al., 2007; Shokouhifar et al., 2019).

The DISC1 protein’s role in neuronal development includes proliferation and migration of the neuronal progenitor cells and synapse formation and maintenance (reviewed in Brandon and Sawa, 2011), and it acts as a molecular hub that interacts with dopaminergic neurotransmission components such as Dopamine (DA) D2 receptors and transporter (Su *et al*., 2014; Trossbach *et al*., 2016; Wang *et al*., 2017). In this regard, the association between DISC1 and the function of DA, one of the leading candidate neurotransmitters in the pathology of different psychiatric disorders, has been profoundly investigated (Davis et al., 1991; Dunlop and Nemeroff, 2007; Cousins, Butts and Young, 2009; Hamilton et al., 2013). The findings suggest that DISC1 has a role in the dysregulation of DA functions, such as the increase in the proportion of striatal D_2_^high^ receptors(Lipina et al., 2010), an increase of DAT levels in the striatum (Jaaro-Peled et al., 2013), or decrease of extracellular DA levels in the nucleus accumbens(Niwa et al., 2010; Dahoun et al., 2017).

A novel transgenic rat model (tgDISC1) has been introduced to study the function of DISC1 in disease and normal cognition. The tgDISC1 rat is a model for aberrant DISC1 signaling by modestly overexpressing non-mutant human DISC1 leading to DISC1 aggregation and thus representing a subset of sporadic cases with mental illness (Leliveld *et al*., 2008). Furthermore, Trossbach *et al*. (2016) reported a full signature of behavioral phenotypes such as amphetamine supersensitivity, hyper-exploratory behavior, and rotarod deficits associated with reductions in DA neurotransmission of the tgDISC1 rats. Thus, this tgDISC1 rat model could be exploited to investigate behavioral differences, specifically variation in reward-related behavior, caused by altered dopamine homeostasis.

Anhedonia, a consequence of deficits in reward processing, is one of the core symptoms of psychotic disorders. It is described as a lack of motivational ability in experiencing pleasure and reduced response to rewarding objects such as non-social reward (e.g., food) or social reward (i.e., social interaction;[American Psychiatric Association, 2012; Thomsen, Whybrow and Kringelbach, 2015]). Considering that the DAergic system acts aas a leading player in the reward learning process, control of motivation (Trifilieff et al., 2013), and encoding the reward prediction error (Schultz, Dayan and Montague, 1997; Cohen *et al*., 2012), anhedonia might be caused by a dysregulation in DA (Chaudhury et al., 2013; Coccurello, 2019).

In addition to general anhedonia, mental diseases such as schizophrenia, depression, autism spectrum disorder (ASD) are characterized by strongly altered social behavior (American Psychiatric Association, 2013). For example, psychiatric patients very often have reduced interest in social interaction and show impairments in social cognition, including deficits in communicative abilities and theory-of-mind (Weightman, Air and Baune, 2014; Green, Horan and Lee, 2015). Since normal social interaction and cognition has been linked to DA activity (Kashtelyan *et al*., 2014; Loureiro *et al*., 2015), the abnormal social behaviors in mental disease may stem from the pathological reward and DA processes, too (Yacubian and Büchel, 2009; Rosenfeld, Lieberman and Jarskog, 2011; Chaudhury *et al*., 2013). However, it is unknown whether general, non-social anhedonia and the social deficits seen in psychiatric disorders stem from the same dopaminergic mechanisms or whether they are the consequence of separate, dissociable processes.

This study exploits the tgDISC1 animal model to address whether abnormal DA homeostasis is linked to reduced non-social reward processing, social interaction seeking, or both. To this end, we compared the choice behavior of tgDISC1 with wild-type rats in a novel paradigm (Seidisarouei *et al*., 2021) in which they chose between the opportunity for social interaction with an unfamiliar juvenile conspecific or drinking sweet solutions with variable sucrose concentrations.

## Methods

### Subjects

The experiment was conducted according to the European Union Directive 2010/63/E.U. for animal experimentation and was approved by the local authorities (Landesamt für Natur, Umwelt und Verbraucherschutz North-Rhine Westphalia, Germany). Transgenic DISC1 (tgDISC1) Sprague Dawley rats and their wild-type (WT) littermate controls were bred at the local animal facility (ZETT, Heinrich-Heine University, Düsseldorf, Germany), 36 Sprague Dawley male rats (tgDISC1 = 12, WT= 12, juvenile rats (WT)=12) in total, consisting of 24 actor rats (PND 57-60, tgDISC1 *M*_*weight*_ = 285 g and WT *M*_*weight*_ = 304 g, at the starting day of the experiment) and 12 juvenile rats (PND 28, *M*_*weight*_ = 145 g at the starting day of the *Social-Sucrose Preference Test (SSPT))*, serving as social stimulus rat. The tgDISC1 actor rats of this study were generated through the identical generation method introduced by (Trossbach *et al*., 2016). Experimental rats were kept in groups of N=2 for actors and N=3 for social stimulus rats, in standard Type IV Macrolon cages in a reversed 12:12 h light-dark cycle. The stable room was kept at a constant temperature of 22°C±2 and a humidity of 55% ±2. Throughout the experiment, all actor rats received standard laboratory rodent food, *ad libitum*, excepting the *Sucrose Discrimination Test* (SDT) phase in which all actors were limited in their food intake (food per rat per day: 22g on weekdays and 25g on weekends).

### Apparatus and behavioral testing

Rats were trained in an X-shaped chambered sociability test (XCST). The apparatus was a radial maze (eight-arm), reduced to a cross/plus-maze setup by removing four arms (Figure 1A), as previously described by (Seidisarouei *et al*., 2021). The maze consisted of a central octagon zone (36 cm diameter, so-called neutral zone) and four arms (60 cm long and14 cm wide) that extended from the neutral zone. Every arm was consistently associated with one specific reward: 3 arms were assigned to three different concentrations of a sucrose solution reward and one arm with a social stimulus rat (see below for details; Figure 1A). During all experimental phases, SDT and Social Sucrose Preference Test (SSPT), only 2 out of 4 arms were kept open, depending on the respective task conditions, to provide a direct preference test between two given rewards. One arm was allocated to the social reward (hereinafter as social arm), and an unfamiliar juvenile rat could be placed on this arm in a fixed cylindrical restrainer built from metal bars and compact plastic for its ceiling and floor (Height: 25.5 cm, Diameter: 17 cm, Ugo Basile Sociability Cage). The restrainer was mounted on the maze at the end of the social’s arm. The social stimulus rat could move around in the restrainer, and the social/physical contact with the actor rat was possible through the openings between the bars. The sucrose solution was provided to the actor rats in a cube plastic dish (8 ⨯ 8 cm) placed at the end of each arm assigned to the non-social reward (i.e., different sucrose concentrations 2%, 5%, and 10%). To facilitate spatial learning of the certain reward in each arm over days, we used sandpaper pieces (17 ⨯ 13 cm) attached to the wall at the entrance of each arm that the actor rats would touch with their whiskers when entering the arms. These sandpapers had varying grades (Figure 1A; 2% sucrose concentration [P800], 5% sucrose concentration [P400], 10% sucrose concentration [P150], and social stimulus [P1200]), following the findings of Guić-Robles, et al. (1989) who have shown that rats, through their whiskers, can differentiate between sandpapers with 200 grains/cm2 and 25 grains/cm2. After each trial, the maze was cleaned by using Ethanol solution (70%).

**Figure 1.**
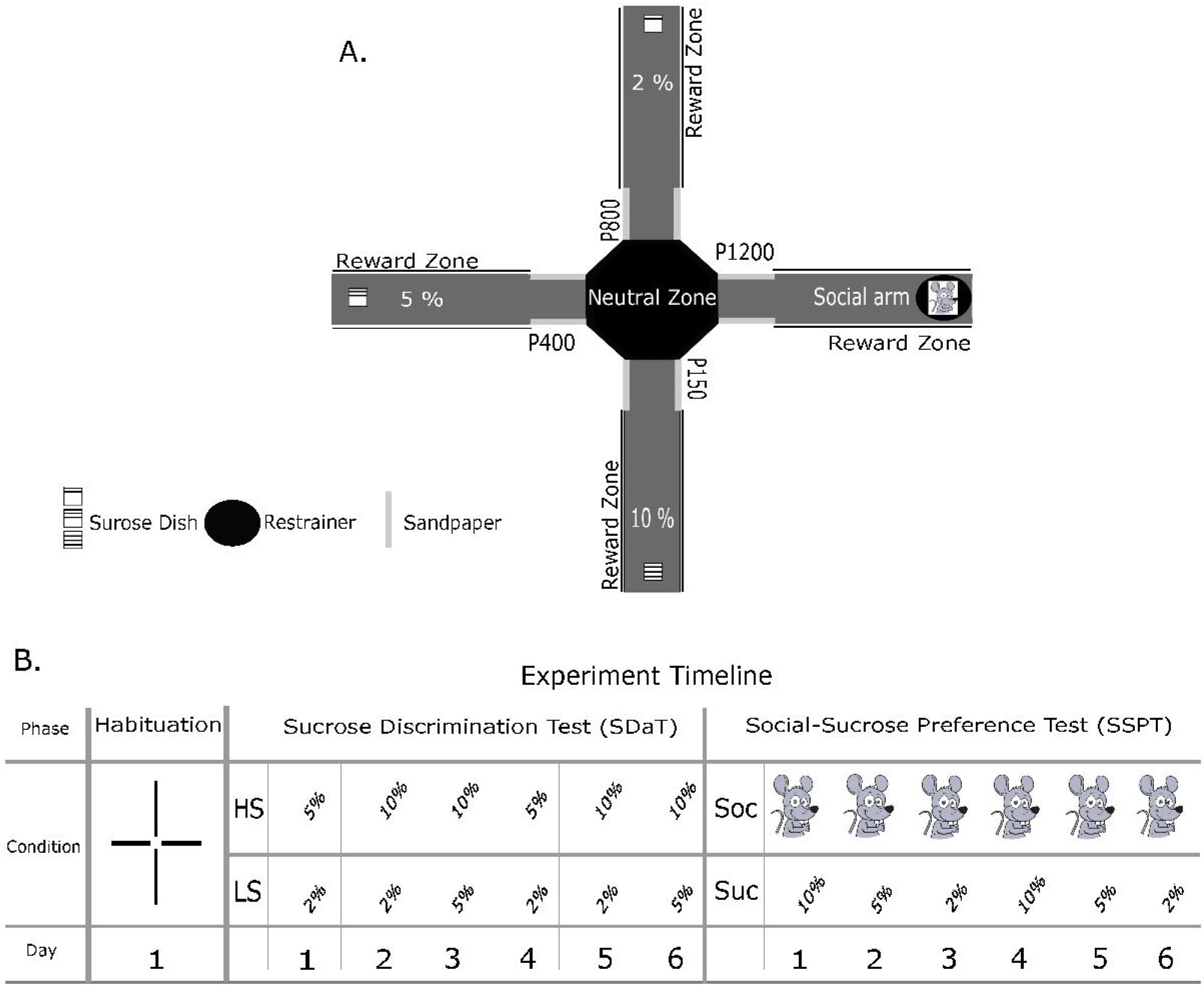
Setup of the study. **A** schematic diagram of XCST maze with non-social reward positions, the restrainer for the social reward, and sandpaper positions and grades. **B** shows an example of the experiment timeline for different phases, days, and conditions. Habituation: free arm investigation in the habituation phase, Sucrose Discrimination Test (SDT): HS; higher sucrose in a given trial, LS; lower sucrose in a trial, Social-Sucrose Preference Test (SSPT): Soc; social reward, and Suc; sucrose.

## Social Sucrose Preference Test

Behavioral testing in the SSPT was divided into three phases: Habituation, SDT, and SSPT (Figure 1B). In all phases, actor rats from both groups (tgDISC1 & WT) were tested daily. In the first phase (Habituation), all four arms were kept open without rewards. Each actor rat explored the maze for 8 minutes. This phase intended to determine whether actor rats were inherently biased towards preferring one given arm or sandpaper grade. The second phase was the SDT which was designed to verify that the rats can indeed distinguish among the three selected sucrose levels (2%, 5%, and 10%). To overcome a potential novelty-induced hypophagia, all actor rats were served 40 ml sucrose a day before starting the SDT phase. Food-restricted actor rats were tested on the SDT phase over six days. On each day, they could choose between two sucrose level concentrations in two repetitions of three different conditions (*2% vs. 5%, 2% vs. 10%*, and *5% vs. 10% in a fixed order*, Figure 1B). In each condition, only two arms were open, and actor rats could freely explore the maze to engage in reward consumption according to their preferences. In each trial, actors were placed in the neutral zone facing not toward open arms at the start of each session (one trial per day). Each trial took 8 minutes; in this time, actors could drink up to 20 ml sucrose solution per plastic dish mounted at the end of each arm. For each new trial and actor, both dishes were filled with fresh sucrose solution. We estimated the time spent in each arm in each condition (see below). After passing the SDT phase, rats were promoted to the SSPT phase. Before starting the SSPT phase, all social stimulus rats were habituated to the experiment room, maze, and restrainer for three days, each day for 8 minutes. To keep baseline motivations equal for both types of reward (social & non-social), after the final day of the SDT, food restriction was lifted to let actor rats gain weight over two days before starting the SSPT phase. For the remainder of the experiment, all rats had access to food *ad libitum*. In the SSPT phase, in each trial with a duration of 8 minutes, the actor rat could freely explore two open arms: the social arm with the unfamiliar social stimulus rat in the restrainer at the end and one of the arms baited with sucrose at the end. There were three conditions in the SSPT (*social reward vs. 2%, social reward vs. 5%, and social reward vs. 10%*). As in the SDT phase, actor rats were tested only once per day; the order of conditions was pseudo-randomized across days and rats (Figure 1B). After completing all conditions, actor rats underwent a second round with the same rat-specific order of conditions as during the first round. Hence, the SSPT phase was completed in six days. Again, we recorded the time spent in each arm on each day of testing as the main index of preference.

In comparison to familiar conspecifics, rats usually spend more time exploring unfamiliar conspecifics (Social Novelty Preference; Smith *et al*., 2015, 2017). Hence, if rats always interact with the same conspecific, the value of social interaction will progressively decline over days with increasing familiarity between rats. To counteract such a trend and maintain the novelty and value of social interaction across testing sessions in the SSPT, twelve different social stimulus rats were used. Thus, each actor rat saw a novel social stimulus rat on each day of testing. The actor-to-social stimulus assignment was counterbalanced across actor rats.

### Behavioral Analysis: Video-Tracking

To track the animals’ position, we used Ethovision (EthoVision XT version 11.5, Noldus). For each phase of the study (Habituation, SDT, SSPT), different tracking arenas were designed. For the phase of Habituation, each arm was divided into two zones (Sandpaper and Reward zone). For the SDT and SSPT, we used the time that the actor rats spent in the reward zones (Figure 1A).

## Data Analysis

In all analyses, the significance level was set at p < 0.05, and all post-hoc tests were Bonferroni-corrected for multiple comparisons. Moreover, the occupancy time for the neutral zone was excluded from all analyses.

### Habituation Phase

To test for spatial bias related to any inherent preference for the different reward and sandpaper zones, we performed two separate two-way repeated-measured ANOVA. The first one assessed the effect of group (tgDISC1/ WT) and sandpaper type (four types) as independent variables (IVs) on the time actors spent in each sandpaper zones as dependent variable (DV), and the second one measured the effect of group (tgDISC1/ WT) and reward zones (four zones) as (IVs) on the time actors spent in each reward zones as (DV). This data was collected during habituation to the maze when rewards were not yet introduced.

### Sucrose Discrimination Performance

To determine whether actor rats discriminated between different sucrose levels in the SDT, we calculated the SDT sucrose solution preference score for each condition and repetition in the SDT as a percentage of time spent in the relatively higher sucrose arm (the arm yielding the higher sucrose concentration on that day; Figure 1B).

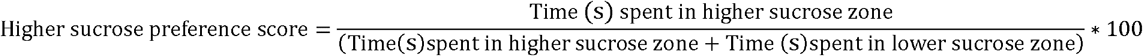

By using these sucrose preference scores, we conducted a three-way mixed ANOVA on the higher sucrose zones preference score (DV) with the group as a between-subject factor (tgDISC1 vs. WT), condition (three levels: *2% vs. 5%, 2% vs. 10%*, and *5% vs. 10%)* and test repetition (first vs. second) as within-subject factors.

To control for potential differences in motor activity between tgDISC1 and WT rats, we additionally measured the distance moved (in cm) per day in the entire maze. We ran an independent samples t-test to analyse the group (tgDISC1/ WT) effect on the distance moved (included all repetitions and conditions).

To explore whether there was a between-group difference in the correlation between *Entrance frequency* and total *duration of stay* (in seconds) in a reward zone, we ran two separate Pearson correlations per group: (1) in the higher sucrose zone. (2) in the lower sucrose zone.

### Social-Sucrose Preference Analysis

for the SSPT, we calculated a social reward preference score:

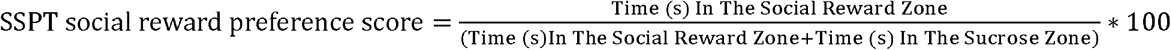

To analyse between-group differences in preference between social (social stimulus rat) or non-social rewards (different sucrose concentrations), we ran a three-way mixed ANOVA to assess the effect of group (tgDISC1 vs. WT), repetition (first vs. second), and condition (*social reward vs. 2%, social reward vs. 5%, social reward vs. 10%*) on the social reward preference score.

Additionally, to determine each group’s preference for social or non-social reward in a given SSPT condition, we conducted six one-sample t-tests versus indifference (50%) on the SSPT social reward reference scores, averaged per animal across the two experimental repetitions.

Again, to determine if there was a difference in the distance moved (cm) between tgDISC1 and WT actors, we conducted an independent samples t-test.

Similar to the SDT, we explored whether there was a between-group difference in the correlation between *Entrance frequency* and total *duration of stay* (in seconds) in a reward zone. We ran two separate Pearson correlations per group: (1) in the social reward zone. (2) in the sucrose zone.

### Software

All statistical analyses were carried out using SPSS Statistics (version 24; IBM, USA), and figures were created using Jupyter Notebook (Kluyver *et al*., 2016) through the packages matplotlib (Hunter, 2007), pandas (McKinney, 2010), ptitprince (Allen *et al*., 2019) and seaborn (Waskom, 2021). For improvement of figures, we used Inkscape (version 0.92.1, Inkscape project, 2020)

## Results

### Habituation Phase

To investigate a potential spatial bias related to any inherent preference for the different reward zones and sandpapers, we executed two distinct repeated measures ANOVAs. These analyses showed no significant bias for either the sandpaper identity or a spatial reward zones location (Table 1)

**Table 1.** Assessment of inherent bias toward a sandpaper or a reward zone. To find out the differences between four sandpaper and also four reward zone we ran two separate two-way mixed ANOVA

|  | Sandpaper Zones |  |  |  | Reward Zones |  |  |  |
| --- | --- | --- | --- | --- | --- | --- | --- | --- |
| | df | F | P-value | $\eta_p^2$ | df | F | P-value | $\eta_p^2$ |
| Group | 1 | 0.105 | 0.752 | 0.009 | 1 | 0.034 | 0.857 | 0.003 |
| Zone | 1.81 | 2.34 | 0.125<sup>a</sup> | 0.176 | 1.67 | 2.81 | 0.093<sup>c</sup> | 0.204 |
| Group x Zone | 1.67 | 0.387 | 0.648<sup>b</sup> | 0.034 | 3 | 0.483 | 0.696 | 0.042 |

a, b, c Greenhouse-Geisser correction.

### Sucrose Discrimination Test

To determine whether actor rats could discriminate between different reward sucrose concentrations (2%, 5%, and 10%), we conducted a three-way mixed ANOVA. Our results did *not* show a significant main effect of group (tgDISC1 vs. WT) on the time spent in the respective higher reward zone (F (1,22) = 0.001, p = 0.978, ANOVA, Figure 2A), but we did find a main effect of condition (2% vs. 5% < 2% vs. 10 and 5% vs. 10%; F(1.57,34.5) = 13.7, p ≤ 0.001, ANOVA, Figure 2B) and repetition (F(1,22) = 29.4, p ≤ 0.001, ANOVA, Figure 2C). Bonferroni-corrected post hoc tests revealed that all rats spent more time in the relatively higher sucrose zone in all three conditions; they spent more time in the 10% zone when the alternative was 5% or 2% sucrose, and they spent more time in the 5% than the 2% zone (Table 2A; Figure 2B). This suggests that all rats were sensitive to relative differences in sucrose concentrations. The Bonferroni test (Table 2B) also showed that rats in both groups significantly spent more time in the higher sucrose zones on the second compared to the first repetition, reflecting learning of the spatial reward arrangement (Figure 2C). The analysis did not reveal any significant interaction effect (Table 3A). Overall, these results demonstrate that all rats learned to express a clear preference order from high to low sucrose. It has been suggested that the dysregulation in dopaminergic signaling in tgDISC1 rats goes along with locomotor hyperactivity (Trossbach et al., 2016). Hence, to ensure that there was no systematic difference in locomotion between tgDISC1 and WT rats in our tasks, we compared the total distance moved across all conditions and repetitions between animal groups. However, we found no significant difference in distance moved between tgDISC1 and WT rats ((t (22) = 0.101, p=0.921, (tgDISC; [M=3536, SE=125] WT; [M=3516, SE=148]), for more details, see Table 2 F&G). Finally, we computed two Pearson correlations per group to determine whether there was a significant correlation between the *frequency of entrance* and *duration of stay* in each zone. The results did not reveal any significant correlations (Table 2 C, Figure 2 D&E).

**Figure 2.**
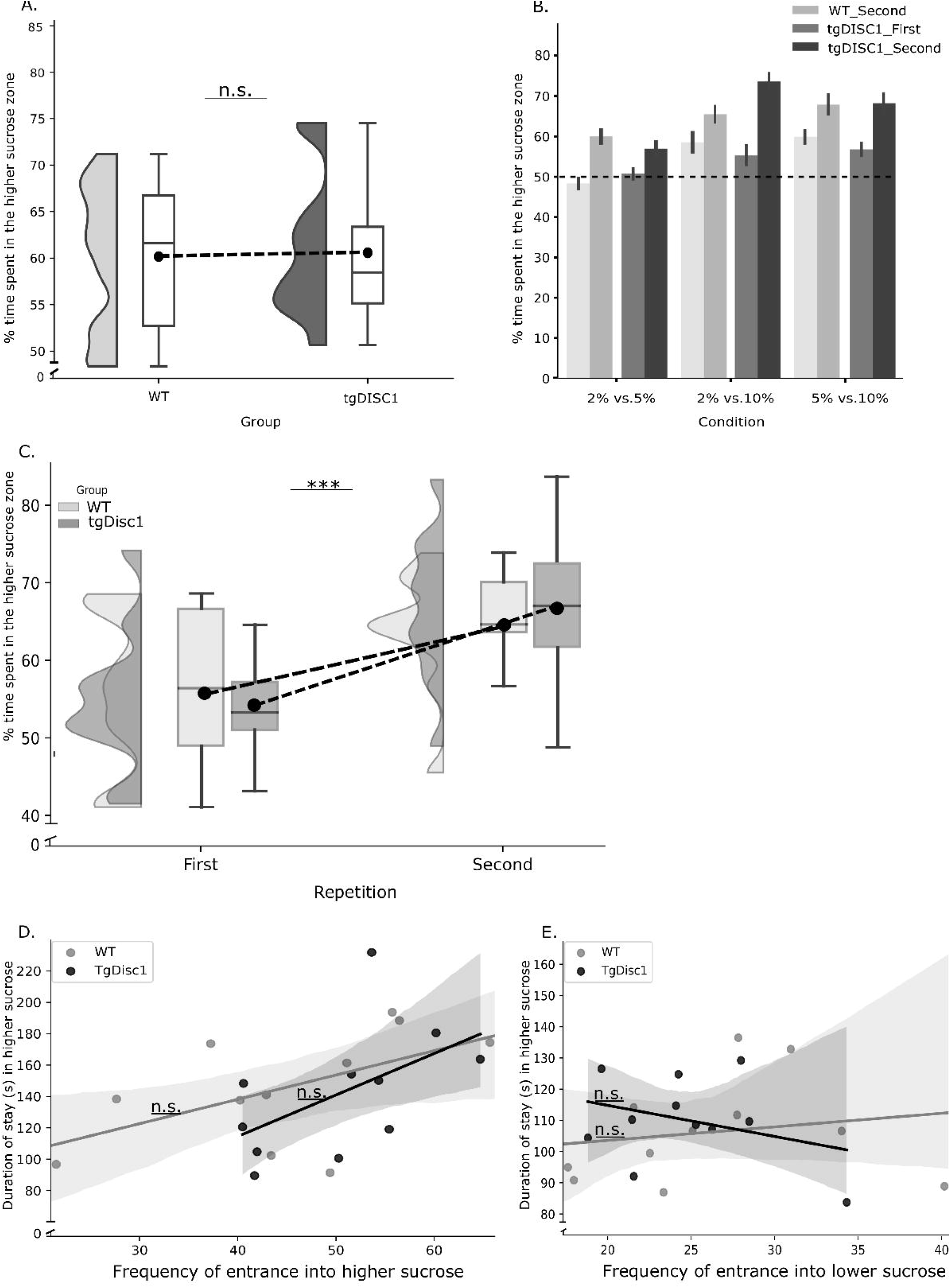
Sucrose Discrimination Test Results. **A** The time spent in higher sucrose zones per group included all conditions and days of SDT. The dashed line connects the mean of the two groups. **B** Time spent in the higher reward zone in each condition; The dashed line indicates the indifference point (50%). **C** The change of time spent in the higher sucrose, across repetitions, per group. The dashed line connects the mean of each group. **D** Correlation between duration of stay and frequency of entrance in higher sucrose zone. **E** Correlation between duration of stay and frequency of entrance in lower sucrose zone. Error bars represent the standard error of the mean (SEM). ***p < .001, n.s.; not significant.

**Table 2.**
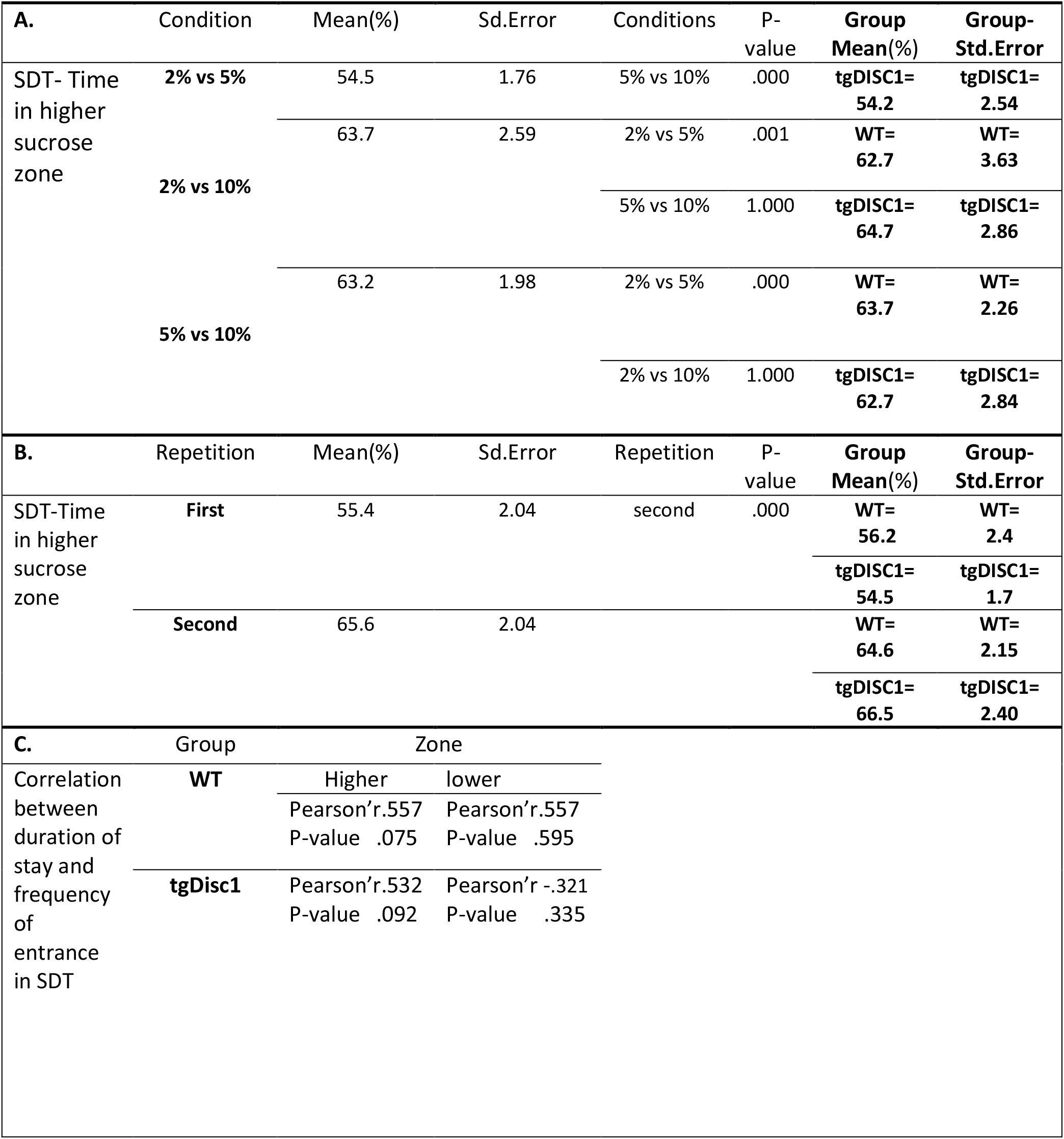
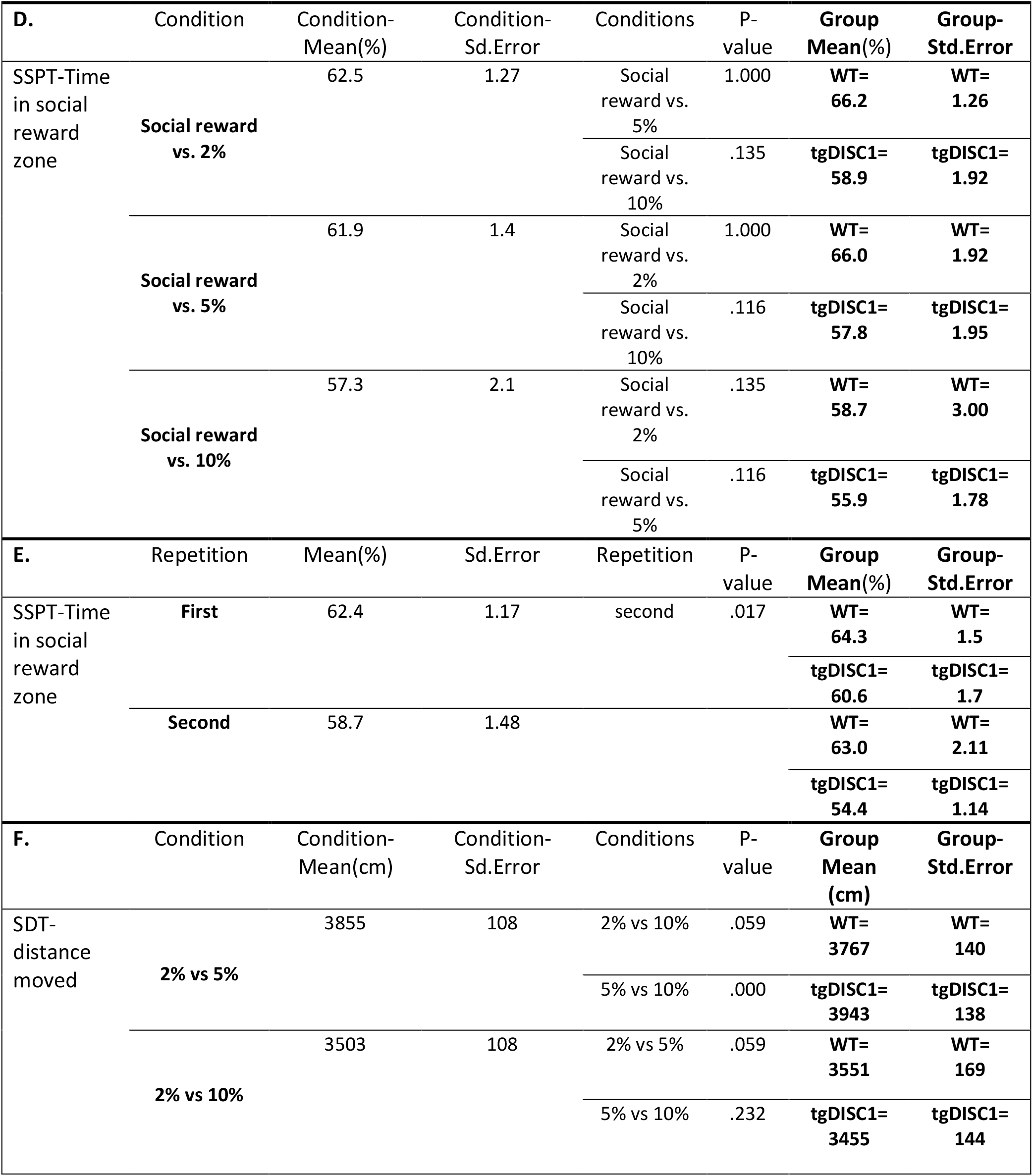
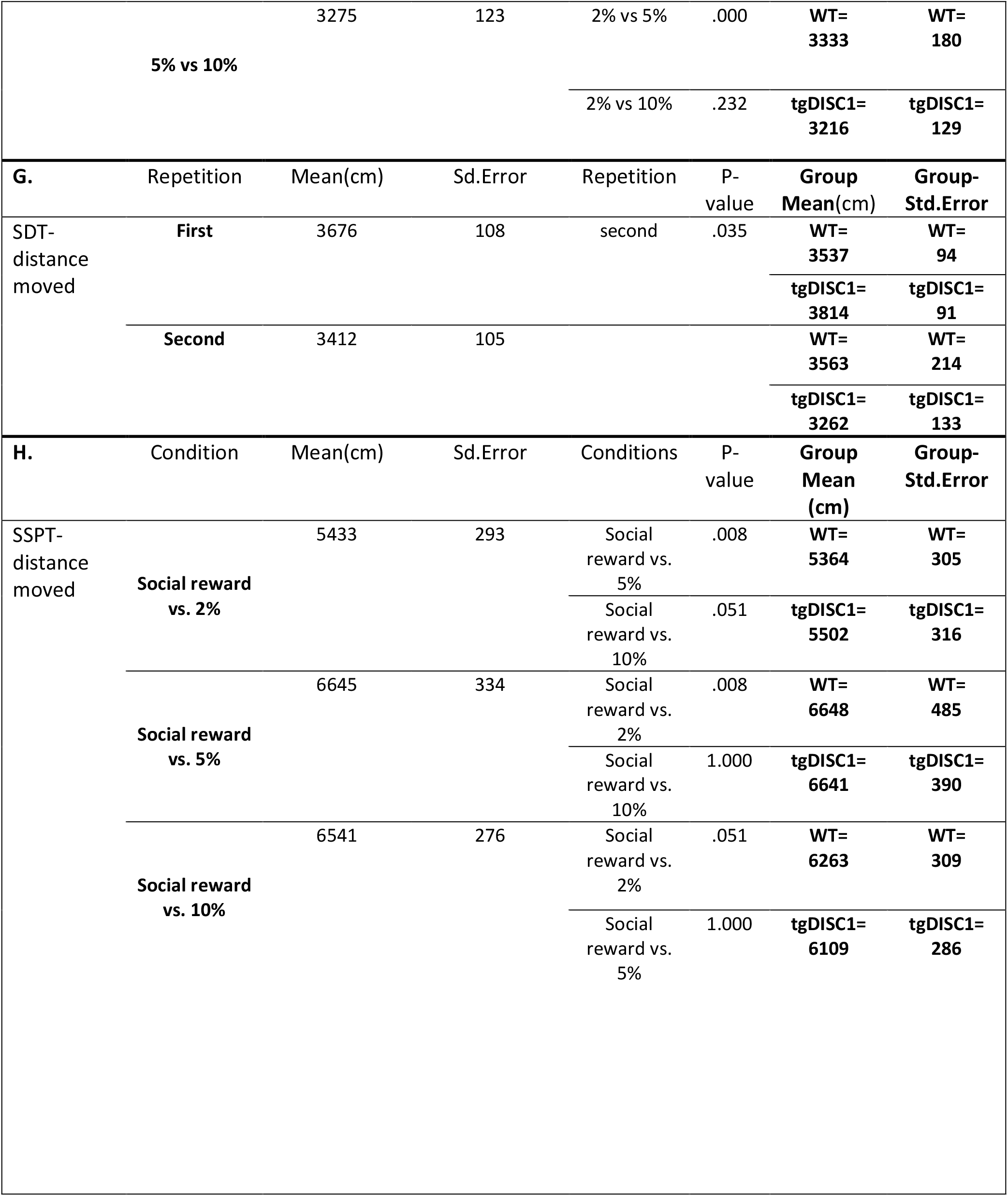
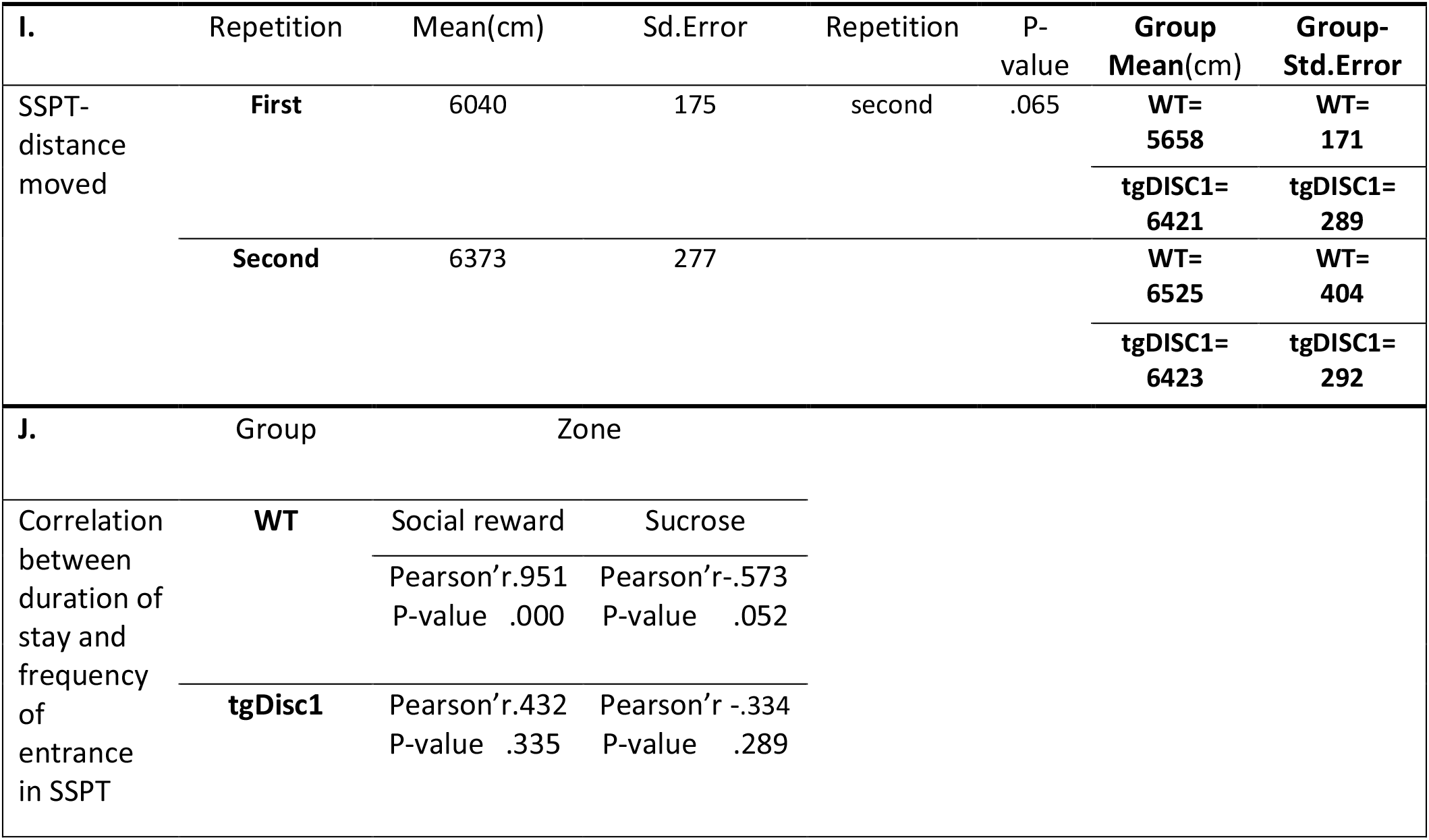
The results of the pairwise comparisons (Bonferroni) for within-subject factors (Condition/Repetition) in both tasks (SDT/SSPT) on IVs (Time in reward zone/Distance moved) respectively and the result of Pearson correlations in both tasks (SDT/SSPT).

**Table 3.**
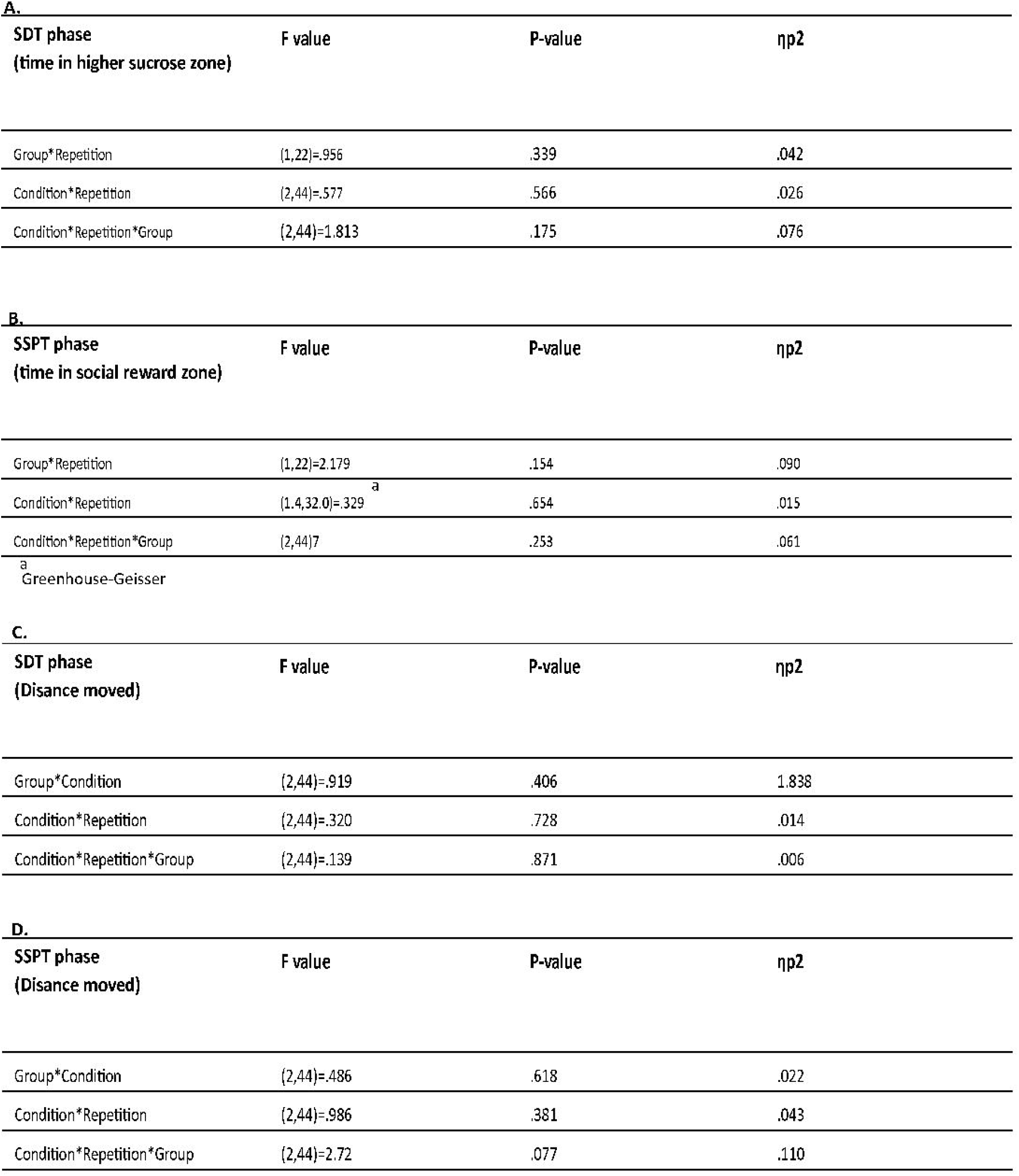
The results of three-way mixed ANOVA (not-significant interaction effects) in both tasks {SDT/SSPT)

### Social Sucrose Preference Test

To investigate between-group differences in the times spent in the respective rewards zones in the SSPT, we ran a three-way mixed ANOVA on the social reward preference score. This analysis revealed a significant difference between groups in the social reward preference score (F (1,22) = 7.3, p =0.013, ANOVA, Figure 3A). This result showed that tgDISC1 rats spent a significantly shorter proportion of time (M = 57.5%, SE = 1.0) in the social reward zone than the WT rats (M = 63.6%, SE = 1.3). There was also a significant main effect of sucrose on the time spent in the social reward zone (F (2,44) = 3.7, p =0.032, ANOVA, Figure 3B). Descriptively, WT rats spent more time in the social reward zone when the alternative was a 2% or a 5% sucrose solution than when the alternative was a 10% solution. However, Bonferroni adjusted post-hoc comparisons (Table 2D) did *not* reveal a significant difference in the percent time rats spent in the social reward zone between the conditions. Accordingly, post-hoc one-sample t-tests against indifference (50%) revealed that all rats spent more time in the social reward zone than in the sucrose zone in all conditions (all t>2.2, p<.05, Fig 3B. I-VI). Next, we also found a significant main effect of repetition on the proportion of time spent in the social reward zone (F (1,22) = 6.6, p = 0.017, ANOVA, Figure 3C). Rats spent significantly more time in the social reward zone on the first than on the second repetition. There were no significant interaction effects. (Table 3B). To make sure that the difference in preference for the social zone between tgDISC1 and WT rats was not the consequence of a general difference in locomotor activity (Trossbach et al., 2016), we, again, compared the total distance moved across all conditions and repetitions between animal groups in the SSPT. However, as in the SDT, we found no significant difference in distance moved between tgDISC1 and WT rats ((t (22) = 0.793, p=0.436, (tgDISC; [M=6372, SE=297] WT; [M=6041, SE=293], for more details, see Table 2 H&I), suggesting that the tgDISC1 effects on social preference are unlikely the result of altered locomotion behavior. Finally, we ran two Pearson correlations per group to determine whether there was a significant correlation between the *frequency of entrance* and *duration of stay* in each zone (sucrose and social rewards). We found a significant correlation between those variables (Table 2J) in the WT rats in the social rewards zone (r (12) = .954, p ≤ 0.001, Figure 3D), indicating that those WT rats that entered the social zone more often also stayed longer. This relationship could not be found in the tgDISC1 rats in the social rewards zone (r (12) = -.432, p = .161, see Table 2J and Figure 3E). Taken together, all rats showed a preference for the social over the sucrose reward in all conditions, but the preference strength, indicated by the percent time interacting with the juvenile, was higher in the WT than the tgDISC1 rats.

**Figure 3.**
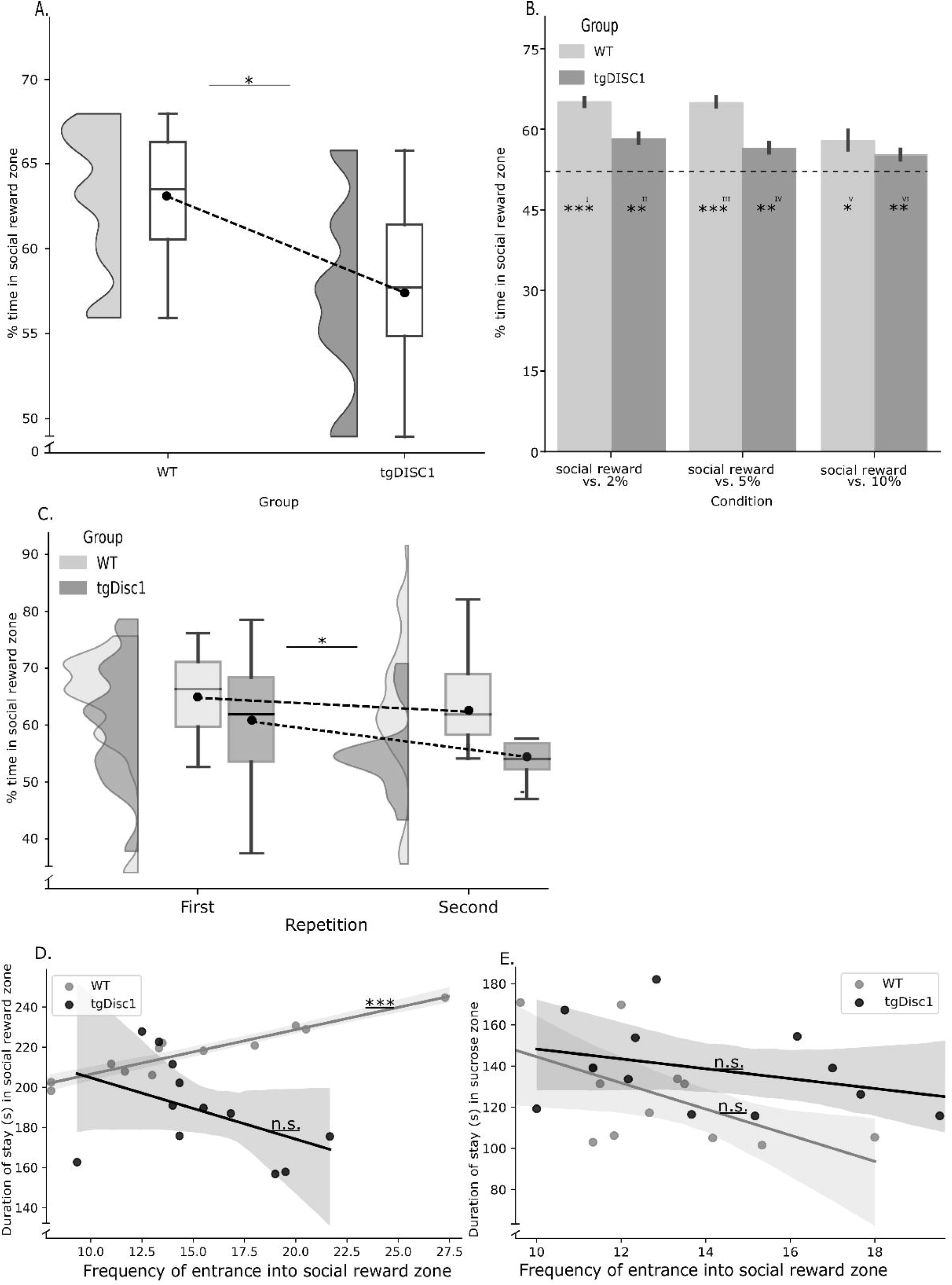
Social Sucrose Preferences Results. **A** between-group difference in percent time spent in the social reward zone (pooled across all days and conditions). The dashed line connects the mean of the two groups. **B** Time spent in the social reward zone. The dashed line indicates the indifference point (50%). **C** Difference in percent time spent in the social reward zone for both groups on the first and second repetition. The dashed line connects the mean of each group. **D** Correlation between duration of stay and frequency of entrance in social reward zone, per group. **E** Correlation between duration of stay and frequency of entrance in sucrose zone, per group. Error bars represent the SEM. *p < .05, **p < .01, ***p < .001, n.s.; not significant.

## Discussion

In this study, we evaluated the effect of tgDISC1 mutations on rat behaviour in a paradigm where the rats could choose between two types of rewards: sweet solutions at variable sucrose concentrations (non-social reward) and the opportunity to interact with a juvenile conspecific (social reward). In our sucrose discrimination task, we found that WT as well as tgDISC1 rats successfully distinguished between sucrose levels, and revealed a clear, well-structured preference for higher sucrose concentrations. Hence, we found no evidence to assume an effect of DISC1 mutation on basic, non-social reward processing. However, when given the choice between drinking sucrose solution or interacting with a conspecific, tgDISC1 rats spent less time with the conspecific than the WT rats, but more time in the non-social reward zones. This might either suggest that, compared to WT rats, tgDISC1 rats had reduced interest in social contact, or that they were lured away from the social interaction zone by the prospect of ingesting more sugar solution in the sucrose zones. However, we consider the latter explanation unlikely since, in the SDT, we found no difference in sucrose preference and sucrose reward seeking behavior between tgDISC1 and WT rats, suggesting that the reduced time that tgDISC1 rats spent with the conspecific in the SSPT was probably not due to hypersensitivity to sucrose rewards, but the result of genuinely reduced interest in social contact.

Why did the tgDISC1 rats attach less value to social interaction? It is plausible to assume that this was the result of altered DA signalling in the brain, in particular in the mesolimbic reward system. Trossbach et al. (2016) reported decreased basal level of DA in striatal samples of tgDISC1 rats. This change was caused by increased D2 receptor and striatal dopamine transporter (DAT) levels, resulting in much faster synaptic DA clearance due to an upregulation of presynaptic DAT. Notably, the upregulation of presynaptic DAT leads to lower net synaptic DA in tgDISC1s. In mice, Hung *et al*. (2017) have shown that an oxytocin-dependent DAergic projection from the VTA to the NAcc Shell region is necessary and sufficient to support real-time social conditioned place preference, strongly suggesting that an altered DA turnover in the NAcc could interfere with social interaction preferences.

Developmentally, previous results (Djouma *et al*., 2006) on the modification (increase in binding) of D2 receptor density by the change in the social environment (social isolation), the role of DAT levels in the regulation of social behaviors (by DAT knockout of mice), (Morgan *et al*., 2002), and the highlighted interplay of regular social contact and striatal function (Fone and Porkess, 2008), all suggest that striatal DA signaling is critical for proper social interactions. Behaviorally, the tgDISC1s rats’ reduced motivation to seek out juvenile conspecifics interaction opportunities aligns with previous studies with neuropsychiatric patients who revealed similar dissocations between social and non-social reward processing (Gotts *et al*., 2012; Lee *et al*., 2019; Butler *et al*., 2020). For example, patients with schizophrenia may experience impairment and disconnection between several components of social motivation required for interactions with positive social outcomes. Likewise, Chevallier et al.(2012) found selective anhedonia (diminished enjoyment) only for social and not non-social reward in children with ASD. Kinard et al. (2020) reported decreased reward prediction error signaling (a critical component of reward-based learning) in frontal brain regions only for social reward in patients with ASD, in line with insensitivity to social rewards found for this group (Dawson *et al*., 1998).

Pinkham et al. (2014) also reported a finding pointing to the distinctiveness of social and non-social information processing in schizophrenia and suggested that individuals with schizophrenia may show a selective impairment in processing social stimuli. Likewise, in depression, Fussner, Mancini, and Luebbe, (2018) found an association between elevated depressive symptoms and decreased approach to social reward (social feedback) but, on the other hand, showed a higher effort by individuals with elevated depression for food rewards.

### Limitations and Future Directions

Rats use different sensory inputs (auditory, olfactory, and visual) in their social interactions. However, it is thought that the most significant rewarding aspect of social interactions for rats is thigmotactic stimulation. In addition, providing a sufficiently large spatial area for social interaction plays an important role in reward experience as well (Kummer *et al*., 2011). In our design, however, rats could only interact through steel bars which potentially decreases the subjectively rewarding experience of the social interactions. Therefore, in future studies, improving the design in a way that facilitates social interactions is recommended.

It has been pointed out that despite activation of the same brain area (ventromedial prefrontal cortex) by both types of rewards (social/non-social), certain areas, such as the amygdala, are more specifically involved in social reward and social cognition (Levy and Glimcher, 2012). Alongside, Sallet et al. (2011) found that the amygdala is involved in perceiving others, and we recently demonstrated that amygdala lesions reduce prosocial behavior in rats(Hernandez-Lallement *et al*., 2016). This possible regional specificity (Ruff and Fehr, 2014) might open up possibilities for local DA transmission reinstatement with the aim of rescuing the DISC1 impairment in social reward processing shown here.

Last but not least, rats communicate through ultrasonic vocalizations (USV) by employing certain call types that are tuned towards social and non-social conditions (Seidisarouei et al., 2021). Therefore, in future studies, in addition to the behavioral/neuronal investigations, recording and analysing USVs in a similar design could shed more light on differences in the subjective affective state of the rats.

## Conclusion

Taken together, the results of this study align with previously found associations between DISC1 and neuronal/behavioural impairments and differences in social vs. non-social reward processing in patients with various psychiatric disorders, suggesting that the tgDISC1 animal model is sensitive to capture the altered social reward processing seen in psychiatric illness, qualifying it as a potential standard for understanding the neural and psychopharmacological basis of abnormal social behavior in mental diseas

## Notes

### Competing Interest Statement

The authors have declared no competing interest.

